# Pangenome comparison of *Bacteroides fragilis* genomospecies unveil genetic diversity and ecological insights

**DOI:** 10.1101/2023.12.20.572674

**Authors:** Renee E. Oles, Marvic Carrillo Terrazas, Luke R. Loomis, Chia-Yun Hsu, Caitlin Tribelhorn, Pedro Belda Ferre, Allison Ea, MacKenzie Bryant, Jocelyn Young, Hannah C. Carrow, William J. Sandborn, Parambir Dulai, Mamata Sivagnanam, David Pride, Rob Knight, Hiutung Chu

**Affiliations:** Department of Pathology, University of California, San Diego, La Jolla, CA; Department of Pediatrics, School of Medicine, University of California, La Jolla, CA; Rady Children’s Hospital, San Diego, CA, United States; Division of Gastroenterology, University of California, San Diego, La Jolla, CA; Center for Microbiome Innovation, University of California, San Diego, La Jolla, CA; Division of Gastroenterology, Northwestern University, Chicago, Illinois; Center for Innovative Phage Applications and Therapeutics (IPATH), University of California, San Diego, La Jolla, CA; Center of Advanced Laboratory Medicine (CALM), University of California, San Diego, La Jolla, CA; Shu Chien-Gene Lay Department of Bioengineering, University of California San Diego, La Jolla, CA; Department of Computer Science & Engineering, University of California, San Diego, La Jolla, CA; Halicioğlu Data Science Institute, University of California, San Diego, La Jolla, CA; Chiba University-UC San Diego Center for Mucosal Immunology, Allergy and Vaccines (cMAV), University of California, San Diego, La Jolla, CA

## Abstract

*Bacteroides fragilis* is a Gram-negative commensal bacterium commonly found in the human colon that differentiates into two genomospecies termed division I and II. We leverage a comprehensive collection of 694 *B. fragilis* whole genome sequences and report differential gene abundance to further support the recent proposal that divisions I and II represent separate species. In division I strains, we identify an increased abundance of genes related to complex carbohydrate degradation, colonization, and host niche occupancy, confirming the role of division I strains as gut commensals. In contrast, division II strains display an increased prevalence of plant cell wall degradation genes and exhibit a distinct geographic distribution, primarily originating from Asian countries, suggesting dietary influences. Notably, division II strains have an increased abundance of genes linked to virulence, survival in toxic conditions, and antimicrobial resistance, consistent with a higher incidence of these strains in bloodstream infections. This study provides new evidence supporting a recent proposal for classifying divisions I and II *B. fragilis* strains as distinct species, and our comparative genomic analysis reveals their niche-specific roles.

## IMPORTANCE

Understanding the distinct functions of microbial species in the gut microbiome is crucial for deciphering their impact on human health. This study reinforces the recent proposal that division II strains constitute a separate species from division I *B. fragilis* strains. Our study provides new evidence that divisions I and II exhibit differential gene abundance related to nutrient utilization, niche occupancy, and virulence. Further, we propose that division I strains are more equipped to colonize the gut and act as commensals, whereas division II strains possess a genetic repertoire for extra-intestinal survival and virulence. Classifying division II strains as *B. fragilis* permits erroneous associations where experimentalists may attribute their findings in division II strains as functions of the better studied *B. fragilis* division I strains. Delineating these divisions as separate species is critical for distinguishing their distinct functions.

## OBSERVATION

*Bacteroides fragilis* is a persistent colonizer of the human gut and has been linked to both health and disease (Wexler, 2007). Multiple studies have reported two distinct, monophyletic groups within *B. fragilis*, referred to as division I and division II, which share 87% average nucleotide identity, while the typical species cutoff is 96% (Johnson, 1978; Podglajen et al., 1995; Ruimy et al., 1996; Gutacker et al., 2000; Nagy et al., 2011; Wallace et al., 2022; English et al., 2023). Here, we use comparative genomics to identify the genetic differences between division I and II strains, to provide further evidence for the classification of these divisions as two distinct species (Wallace et al., 2022; English et al., 2023). We examined genes conserved within each division, but not between divisions, which likely play a fundamental role in their biology and function within their respective niches. This comprehensive analysis not only enhances our understanding of *B. fragilis* but also provides valuable insights into the properties and functions of division I and II strains and their contribution to host-microbe interactions.

We analyzed a total of 694 whole genome sequences, 139 from our own collection, which we isolated and sequenced for the first time (Sanders et al., 2019), and the remaining from public sources (**Table 1 and 2**). To compare the genetic relatedness between divisions, we employed MASH, a whole genome k-mer-based approach (Ondov et al., 2016) to determine the genetic distance between each strain (**Figure 1A**). Metric multidimensional scaling (mMDS), which visualizes the pairwise dissimilarities or distances between a set of objects in a lower-dimensional space, automatically revealed a clear separation of strains into two distinct divisions (**Figure 1A**). To further support this distinction, we found a significant difference in GC content (p=8.1e-5) **(Figure 1B)**, though no differences in genome size (p = 0.22) **(Figure 1C)**. Based on the phylogeny of the core genome alignment by maximum likelihood, midpoint-rooted, divisions I and II also separate into discrete clades **(Figure 1F)**. Collectively, these analyses reinforce the recent proposal to classify *B. fragilis* division II strains as a novel species (Wallace et al., 2022; English et al., 2023).

**Figure 1:**
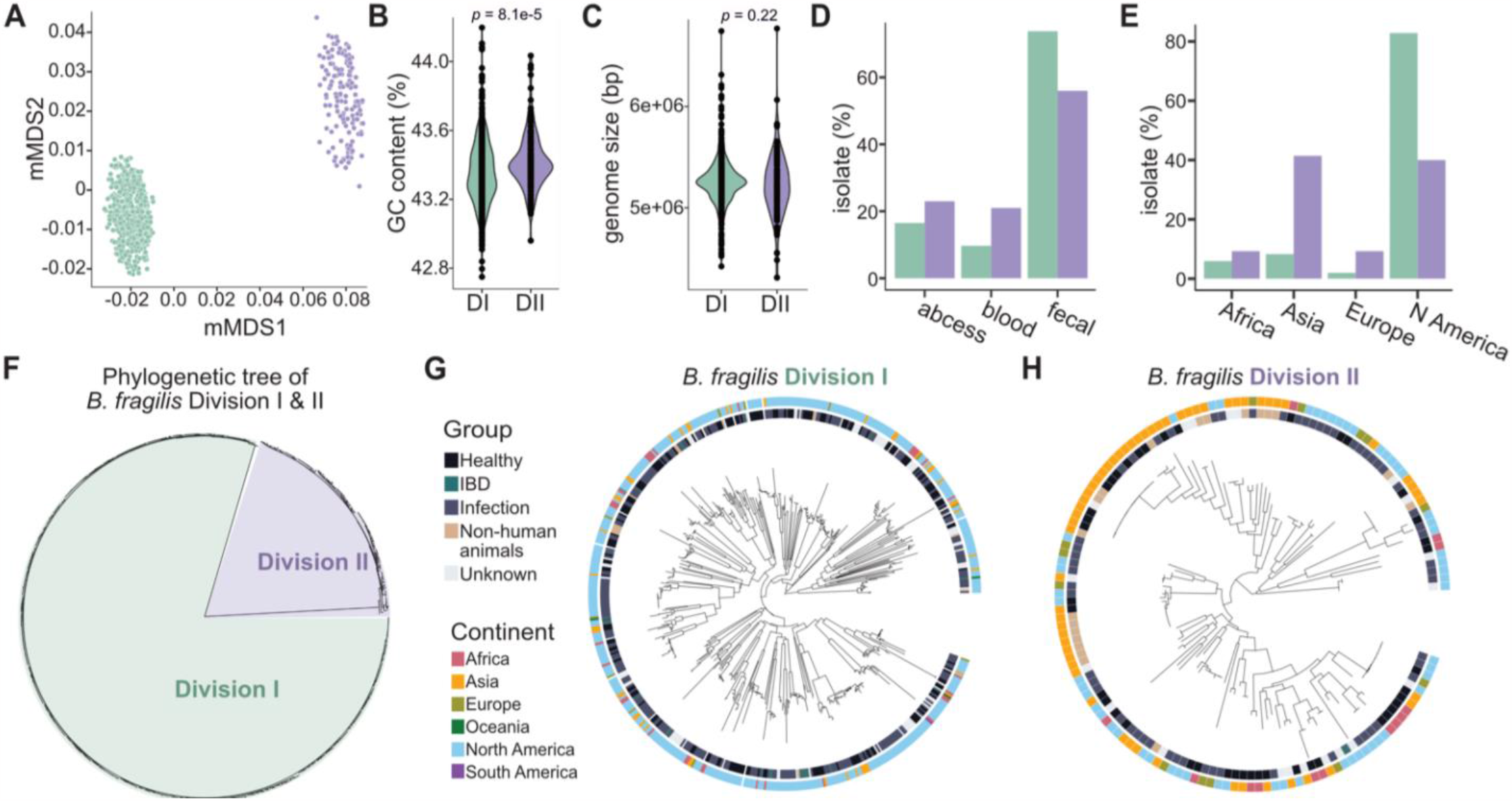
*B. fragilis* is composed of two monophyletic divisions. A) Metric multidimensional scaling (mMDS) of the k-mer based MASH distances of 694 strains, colored by division I (green, n=554) and II (purple, n=140). B) GC content (%) of isolate assemblies in division I and II isolates. Average for division I = 43.35% and division II = 43.42% (p = 8.1e-5, Welch’s t-test with unequal variance; n=694). C) Genome size (bp) of isolate assemblies in division I and II isolates. Average for division I=5.26 x 10^6^ bp and division II=5.22 x 10^6^ bp (p = 0.22, Welch’s t-test with unequal variance; n=694). D) The proportion of isolates originating from abscess (p=0.18), blood (p=0.0049), and fecal (p=0.0011) samples in division I (green) compared with division II (purple), p-values from Fisher’s Exact Test. Division I: n=309, fecal=228, blood=30, abscess=51; Division II: n=100, fecal=56, blood=21, abscess=23. E) The proportion of isolates originating from Africa (p=0.18), Asia (p= 2.2e-16), Europe (p=0.00019), or North America (p= 2.2e-16) in division I (green) compared with division II (purple), p-values from Fisher’s Exact Test. Division I: n=554, Africa=33, Asia=46, Europe=11, North America=459; Division II: n=140, Africa=13, Asia=58, Europe=13, North America=56. F) Phylogenetic tree of the core genome alignment of 638 strains through maximum likelihood, midpoint rooted, colored by division I (green) and II (purple). G) The phylogenetic tree of the core genome alignment of division I strains through maximum likelihood, midpoint rooted, annotated with the inner ring, Group: healthy, infection, IBD, non-human animal, unknown; and outer ring, Continent: Asia, Africa, Europe, Oceania, North America, South America (n=554). H) The phylogenetic tree of the core genome alignment of division II strains through maximum likelihood, midpoint rooted, annotated with the inner ring, Group: healthy, infection, IBD, non-human animal, unknown; and outer ring, Continent: Asia, Africa, Europe, Oceania, North America, South America (n=140).

We next investigated whether divisions I and II associate with different disease states, isolation sites, or other metadata categories. In our survey of 694 strains, we found division I strains comprised 80% of the total (554 of 694). Among the 409 strains isolated from abscesses, fecal samples, or blood, 74% of division I strains originated from fecal samples, compared with 56% of division II (p=0.0011) (**Figure 1D**). Additionally, 16% and 10% of division I strains were isolated from abscesses or blood, respectively, compared to 23% and 21% of division II strains (blood, p=0.0049; abscess, p=0.18) (**Figure 1D**). Notably, division I and II strains exhibited variations in the continent of isolation. 80% of division I strains originated from North America, compared with only 40% of division II strains (p=2.2e-16) (**Figure 1E, H**). In contrast, only 8% of division I strains originated from Asia, compared to 41% of division II strains (p=2.2e-16) (**Figure 1E, G**). To further explore the geographical distribution of these divisions, we examined 502 species-genome bins (SGBs) classified as *B. fragilis*, which were reconstructed from 9,428 metagenomic samples worldwide (Pasolli et al., 2019). The results revealed that 437 strains belonged to division I, whereas 65 were division II. No sample contained both divisions, in line with reports from other studies (Rashidan et al., 2018). Most of the division I strains (75%) originated from Europe or North America, whereas most division II strains (60%) were from Asia. This aligns with previous reports indicating a higher rate of *cfiA*+ isolates (division II) in Japan, Hong Kong, and India (Cao et al., 2022). Altogether, division II strains are more prevalent in Asian countries compared to Western populations, and the under-representation of division II strains in public strain repositories may be a result of under-representation of specific populations (Abdill et al., 2022).

Our data further support the idea that divisions I and II represent distinct genomospecies. Therefore, we next tested whether these divisions exhibit differing metabolic requirements, ecological niches, or lifestyles. We compared the pangenomes of the *B. fragilis* divisions using panpiper (*rolesucsd/Panpiper*, n.d.), and identified 794 differentially prevalent genes (log-fold change ≥ 2) (**Figures 2A-B and Table 3**). Each of the *B. fragilis* divisions exclusively harbored either the *cfiA* (division II) or *cepA* (division I) gene (**Figure 2E and Table 3**), as previously described (Parker & Smith, 1993; Rasmussen et al., 1990). We then assessed the differential abundance of carbohydrate-active enzymes, along with reference metabolic (EC) and reference KEGG orthology pathways (KEGG KO) (**Figures 2C-E**). Within division II strains, all upregulated glycosyl hydrolase (GH) categories (GH5, GH9, GH51, and GH95) are associated with the degradation of plant cell walls (**Figure 2C**). Specifically, BFAG_03498 (ko:K01179, GH9) is predicted to mediate the breakdown of cellulose (Béguin, 1990), BFAG_02344 (GH51) is involved in the breakdown of arabinose-containing polysaccharides, and BFAG_0465 (GH95), an alpha-L-fucosidase, cleaves internal beta-1,4-glycosidic bonds which are common in seaweed and mushrooms (Wu et al., 2023) (**Table 3**). One possible explanation for an increased abundance in plant cell wall degradation genes in division II strains is differences in diet between hosts harboring division I versus II strains, which could correlate with their differential geographic abundance (De Angelis et al., 2020). In contrast, in division I strains, we identified several genes and pathways associated with the degradation of complex carbohydrates, a hallmark feature of gut-resident commensal *Bacteroides* (Wexler, 2007) (Pudlo et al., 2022). Specifically, we identified two predicted alpha-L-rhamnosidases (GH78; BF9343_0522, BF9343_0310), which are core genes exclusive to division I (**Figures 2C and Table 3**). Because humans cannot cleave terminal rhamnose units, rhamnosidases play an important symbiotic role, releasing rhamnose in the human gut, which can then be converted into the short-chain fatty acid propionate (Mueller et al., 2018).

**Figure 2:**
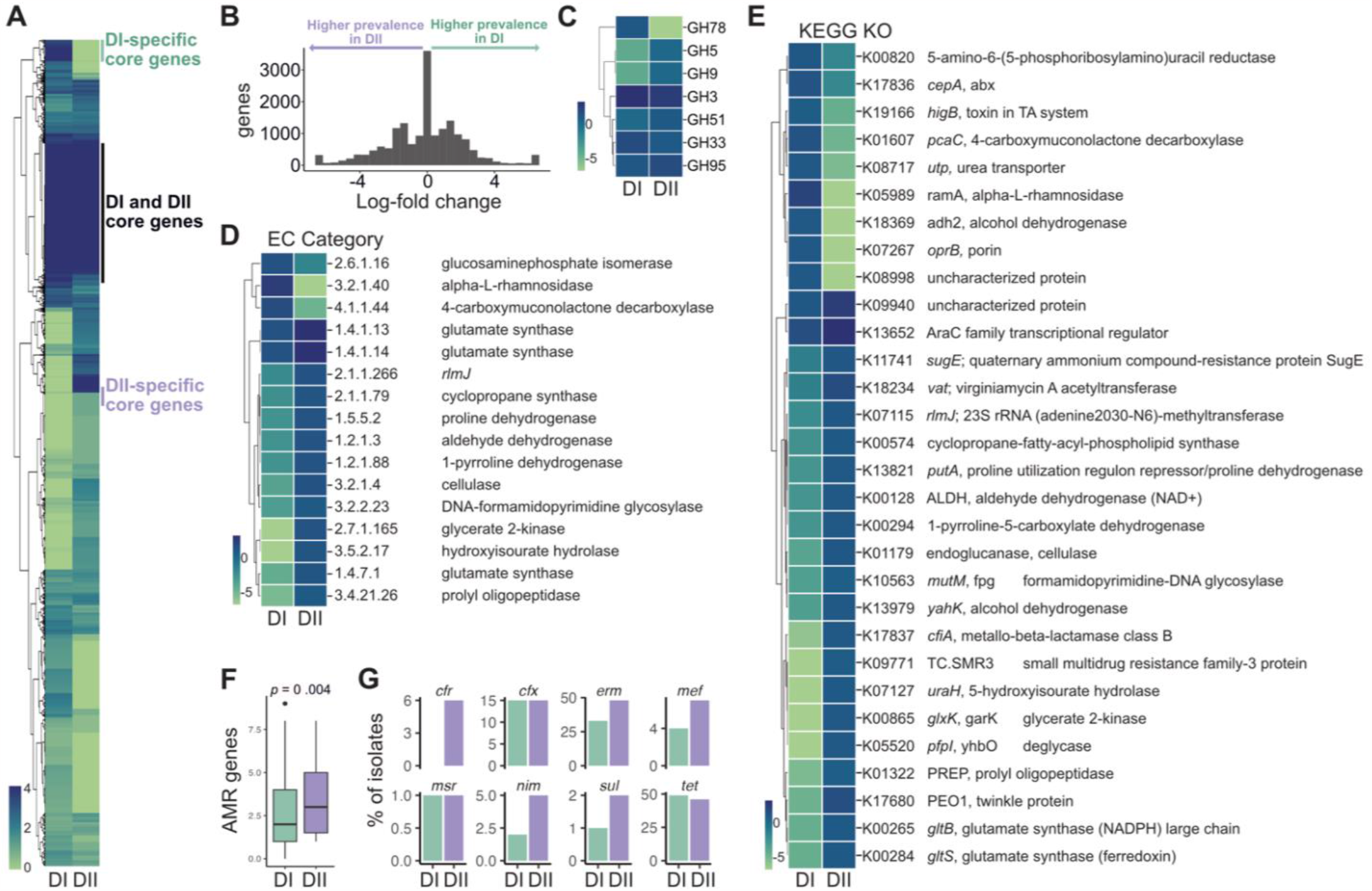
*B. fragilis* Divisions I and II segregate by multiple differentially abundant genes and gene categories. A) Relative log gene abundance heatmap summarized by division, where genes are clustered by R pheatmap complete method, annotated by regions of gene clusters core to both divisions, core only to division I, or core only to division II. B) Histogram of log_2_-fold change of prevalence between all genes in division I versus II. C-E) Log_2_ average number of genes per isolate in categories C) Carbohydrate-Active Enzymes (CAZy) (log-fold change ≥ 0.5), D) EC category (log-fold change ≥ 1), and E) KEGG KO (log-fold change ≥ 0.5) between divisions I and II, displaying categories significant by Kruskal-Wallis test (corrected p ≤ 0.01). Legend is log_2_ average number of genes per isolate in each category. F) Total number of antimicrobial resistance (AMR) genes per isolate for each division divisions, p = 0.004, Welch’s t-test. G)The percentage of isolates per division with each antimicrobial resistance gene.

Division I strains also exhibit an enrichment of GH33 sialidases, which catalyze the cleavage of terminal sialic acid residues (**Figure 2C**). While sialidases have been linked to virulence (Godoy et al., 1993), our previous work established a role for *B. fragilis* GH33/NanH sialidase in intestinal colonization and persistence during early life (Buzun et al., 2023). Furthermore, as sialic acid is identified in capsular polysaccharides and lipooligosaccharides (Ghosh, 2020), its presence may influence colonization and interactions within the host. Additionally, the type VI secretion system GA3 (T6SSiii) is more abundant in division I strains (86%) compared to division II (39%). This system, exclusive to *B. fragilis*, is recognized for mediating intra-strain competition and influencing colonization dynamics (Sheahan et al., 2023). Thus, the differential abundance of GH33 sialidases and T6SSiii GA3 suggests distinct colonization strategies within the gut.

Division II strains may play a different role in niche occupancy, with several differentially prevalent genes correlated with pathogenicity. Notably, division II strains exhibit an increased abundance in genes related to proline degradation and glutamate synthesis pathways (**Figure 2D and Table 3**), known for their association with virulence in several bacterial species (Krishnan et al., 2008; Nakada et al., 2002; Zheng et al., 2018). Prolyl oligopeptidase (EC 3.4.21.26; BFAG_03703) initiates proline cleavage from short peptides, leading to subsequent degradation of free proline by PutA (EC 1.5.5.2; BFAG_03859), which oxidizes proline to glutamate and serves as a transcriptional regulator for essential virulence factors (Moxley et al., 2011; Ye et al., 2022). Proline catabolism, linked to colonization, persistence, and protection from stress, including oxidative and osmotic stress, has been associated with the virulence of several bacterial species (Nakada et al., 2002; Zheng et al., 2018). The higher abundance of multiple genes linked to proline degradation in division II strains suggests their potential to effectively respond to oxidative stress and adapt to extra-intestinal niches, supporting their association with bloodstream infections (Jeverica et al., 2019). Moreover, division II strains have an increased abundance DNA-formamidopyrimidine glycosylase (EC 3.2.2.23; BFAG_03121), which plays a crucial role in processes leading to recovery from mutagenesis and/or cell death caused by alkylating agents (**Figure 2D, Table 3**). These adaptive mechanisms may confer a survival advantage to division II strains in specific environments.

Finally, we observed differential prevalence in genes and pathways related to multidrug resistance. Within division I, we identified an increased prevalence of gamma-carboxymuconolactone decarboxylase (EC 4.1.1.44) (**Figure 2D**), implicated in the degradation of aromatic compounds and associated with antimicrobial resistance (AMR) (Rana et al., 2023). We identified a putative erythromycin esterase that detoxifies macrolides also more abundant in division I (Zieliñski et al., 2021). In contrast, division II strains have a higher abundance of efflux proteins (K09771, K11741) (**Figure 2E and Table 3**). Additionally, virginiamycin A acetyltransferase (*vat*, K18234), providing resistance to streptogramins, is more prevalent in division II (**Figure 2E and Table 3**). Division II strains harbor a higher number of known antimicrobial resistance genes per isolate compared with division I (p = 0.004) (**Figures 2F and 2G**). Collectively, these findings suggest that division II may have a higher potential for virulence compared to division I strains. Further characterization of the functional impact of the genes unique to each division is essential for understanding their roles and interactions within the intestinal ecosystem and host.

Altogether, our comprehensive analysis revealed distinct genetic profiles and functional pathways that differentiate *B. fragilis* divisions. The pangenome of division I strains aligns with their recognized role as commensals and proficient gut colonizers in the mammalian host. Conversely, division II strains harbor a unique collection of genes associated with plant cell wall degradation, suggesting a correlation with their higher abundance in Asian countries or dietary preferences. The presence of genes mediating survival in toxic environments highlights the adaptive capabilities of division II strains. Importantly, these genetic distinctions may underlie the higher prevalence of division II strains in bloodstream infections. Collectively, our comparative genomics study unveils distinct genetic signatures within *B. fragilis* divisions, offering insights into their intricate interactions with the host and respective ecological niches.

## Acknowledgements

We thank members of the Chu lab for technical support and helpful discussions. This work was supported by grants from the National Institute of Health (NIH) R01 AI167860 and P30 DK120515. Additional support was provided to H.C. by the Chiba University-UC San Diego Center for Mucosal Immunology, Allergy and Vaccines (cMAV), CIFAR Humans and the Microbiome Program, and The Hartwell Foundation. Support to R.E.O. was provided by T32 AR064194 (NIAMS). Support to M.C.T was provided by T32 DK007202 (NIDDK), the National Academies of Sciences, Engineering and Medicine through the Predoctoral Fellowship of the Ford Foundation, and the Howard Hughes Medical Institute (HHMI) Graduate Fellowships grant (GT15123). This publication includes data generated at the UC San Diego IGM Genomics Center utilizing an Illumina NovaSeq 6000 that was purchased with funding from a National Institutes of Health SIG grant (S10 OD026929).

